# Diversity-generating retroelements enable phages to expand their host range in the human gut

**DOI:** 10.64898/2026.10.30.763091

**Authors:** Anne W. M. van den Hurk, Michele Zuppi, M. David Kuida, Max Nieuwdorp, Koen Wortelboer, Hilde Herrema, Patrick A. de Jonge

**Author notes:** These authors contributed equally as senior authors (Hilde Herrema, Patrick A. de Jonge).

## Abstract

Diversity-generating retroelements (DGRs) are genetic elements that use error-prone reverse transcription to hypermutate specific target genes, most often the receptor-binding proteins of phages. They are enriched in human gut viruses, where they are hypothesised to enable phages expand their host range. Yet DGR-mediated host-range expansion across bacterial species has so far only been predicted *in silico* and has not been examined directly within complex microbial communities. To test this hypothesis *in situ*, we tracked phage-host interactions in the human gut, combining bulk metagenomic, virus-like particle (VLP), and Hi-C sequencing of longitudinal faecal samples collected over a five-week period. This culture-independent approach identified 380 interactions between 97 phages and 76 bacterial hosts. Nearly a third of these phages (31 of 97) encoded a DGR, and DGR-encoding phages were more likely to interact with multiple bacterial species. Moreover, nucleotide diversity in DGR-targeted genes increased with the number of hosts a phage interacted with and was concentrated in the surface-exposed, DGR-targeted variable region of its receptor-binding proteins. Together, these results provide *in situ* evidence that DGRs enable phages to expand their host range in the human gut.

## Introduction

Bacteriophages (phages) are viruses that infect bacteria and are present in every known ecosystem, including the human gut microbiome^1,2^. Through lysis, lysogeny, and horizontal gene transfer, phages shape the composition and functioning of bacterial communities. This makes phages promising candidates for controlling such communities, such as in food-safety applications or as phage therapy^3,4^. To do so, however, it is necessary to understand which bacteria a given phage can infect, i.e. its host range.

Phage host ranges have long been considered narrow, typically spanning no more than a single bacterial species, or even one strain. Yet it is becoming increasingly clear that broad-host-range phages infecting multiple bacterial species are common across ecosystems^5–7^. Host ranges can also shift over time, shaped by phenotypic variability as well as ongoing co-evolution between phages and bacteria^8^.

Phages use several strategies to achieve a broad host range. Some phages are polyvalent, expressing multiple receptor-binding proteins simultaneously that each recognise a different bacterial host^9–11^. Others engage in host switching: while an individual phage particle can infect only one host, mutations allow the phage species as a whole to infect multiple bacterial hosts over time^7^.

A distinctive system that enables host switching is the diversity-generating retroelement (DGR)^12,13^. DGRs use an error-prone reverse transcriptase to generate cDNA of a template region (TR), which is then inserted into a homologous variable region (VR) within specific target genes. Because of the error-prone, adenine-specific nature of this reverse transcription, it introduces extensive mutations in the VR of the target genes, which often encode a receptor-binding protein^12^. DGRs are therefore hypothesised to enable phages to adapt to their bacterial host, and potentially to expand their host range in the process^13^. To date, however, DGR-mediated expansion of phage host ranges across bacterial species has only been predicted *in silico*^13–15^, and has not been examined directly within complex microbial communities.

Given that DGRs are enriched in the human gut^13^, this environment is well suited to examine the host range of DGR-encoding phages *in situ* within a complex microbial community. Here, we set out to test whether DGRs enable phages to interact with multiple bacterial species. To this end, we analysed longitudinal human faecal samples using bulk metagenomic, virus-like particle (VLP), and Hi-C sequencing. Hi-C sequencing is a form of proximity ligation sequencing that crosslinks and sequences physically adjacent DNA. When a phage infects a bacterial cell, the phage and bacterial DNA become co-localised and are crosslinked together, allowing phage-host interactions to be inferred in a culture-independent manner. Using this approach, we tracked the dynamics phage-host interactions over time and found that DGRs are enriched in phages interacting with multiple bacterial hosts. Moreover, the genes targeted by DGRs showed increased nucleotide diversity specifically when phages interacted with multiple hosts, providing direct support for the hypothesis that DGRs enable phages to expand their host range in the human gut.

## Methods

This study was previously published^16^ and was approved by the Medical Research Ethics Committee Amsterdam UMC (METC 2018_231). The study was executed in line with declaration of Helsinki principles and all participants signed written informed consent before taking part. The study was registered at the Dutch National Trial Register (NTR) under NL8289.

### Bulk metagenomic and Hi-C sequencing

Bulk metagenomic and Hi-C sequencing were done by Phase Genomics. Hi-C libraries were prepared with the ProxiMeta™ Hi-C Kit (Phase Genomics, Protocol v3.0) following the manufacturer’s protocol. Briefly, 250 mg of faeces was homogenized and pelleted in PBS. Samples were incubated for 20 min in crosslinking solution at room temperature, after which the crosslinking was stopped. The cells were then lysed by bead beating. The chromatin was pelleted and bound to beads, before being fragmented, end-filled with biotinylated nucleotides, and proximity-ligated. After crosslink reversal, the DNA was purified and biotinylated ligation junctions were enriched with streptavidin beads. Dual unique-indexed, Illumina-compatible libraries were then constructed from the bead-bound DNA. For bulk metagenomic sequencing, DNA was extracted with the ZymoBIOMICS DNA Miniprep Kit (Zymo Research). Libraries were prepared from 50 ng of input DNA using the Nextera (Illumina).

### VLP sequencing

VLP sequencing was also done as previously described^16^. Briefly, VLPs were extracted from 500 mg of faeces using high-speed centrifugation and filtration through a 0.45 µm membrane. Free-DNA debris was digested, VLPs were lysed, the DNA was purified using a two-step phenol/chloroform extraction protocol, and further purified using the DNeasy Blood & Tissue kit (Qiagen Cat#69506). Library preparation was done with the NEBNext Ultra II FS DNA library prep kit (New England Biolabs Cat#E7805L) and the NEBNext Multiplex Oligos for Illumina dual indexes (New England Biolabs Cat#E7600S). Quality and concentration of the VLP libraries were assessed with the Qubit dsDNA HS kit (ThermoFisher Cat#Q32854) and the Agilent High Sensitivity D5000 ScreenTape system (Agilent Technologies). Libraries were sequenced using 2×150 bp paired-end chemistry on an Illumina NovaSeq 6000 platform with the S4 Reagent Kit v1.5, 300 cycles (Illumina Cat#20028312) at the Core Facility Genomics of the Amsterdam UMC. To check for contamination, negative controls were included DNA extraction and library preparation, which did not yield any measurable DNA and were therefore not sequenced.

### Read quality control and assembly

All software was used with standard settings unless noted otherwise. VLP and bulk metagenomic sequencing reads were quality controlled by trimming with fastp^17–19^ v1.1.0 (option -- detect_adapter_for_pe), error correction with tadpole from bbmap^20^ v39.01 (options mode=correct, ecc=t, prefilter=2). High-quality reads were assembled into contigs using MEGAHIT^21,22^ v1.2.9, which were filtered for sequences longer than 2,500 bp for downstream analysis. This yielded one assembly per timepoint for the bulk metagenomic and another for the VLP sequencing data. Contigs were then concatenated per person, and identical copies were removed with dedupe from bbmap v39.01 (minidentity=100).

### Viral sequence identification

To determine which sequences were viral, the per-sample assemblies were run through VirSorter2^23^ v 2.2.4, after which phage sequences from the different days and data types were concatenated per person. After removing duplicates with SeqKit^24^ v2.10.0 (rmdup), CheckV^25^ v1.0.1 contamination was used to determine the edges of prophage regions, which were then trimmed off. Trimmed viral contigs were then clustered into viral species at 95% ANI over 85% length with Vclust^26^ v 1.3.1, after which the longest sequence per viral species was selected as its representative. Species representatives were annotated with Pharokka^27^ v1.9.1 and Phold^28^ v1.2.6, and their completeness was determined with CheckV^25^ v1.0.1. DGRs were identified using scripts from Roux *et al.*^13^, which, briefly, identify reverse transcriptases based on HMM profiles using hmmsearch^29^, detect repeats around the candidate reverse transcriptases, and select putative DGRs based on the repeat patterns and reverse transcriptases length^13^. DGR clades were assigned by adding the reverse transcriptase sequences of our identified DGRs to the reference alignment of Roux *et al.* with MAFFT^30^ v7.526 (--add --keeplength --genafpair -- maxiterate 1000). Branch lengths and model parameters of the reference were optimized using RAxML-NG^31^ v. 2.0.2 (--evaluate --model WAG+G). Our reverse transcriptases were then placed on the fixed reference tree using EPA-ng^32^ v0.3.8 and grafted with gappa^33^ v0.9.0 (examine graft) and assigned to the reference clade they were placed in.

### Host binning

Contigs were binned with MetaBAT2^34^ v2.15 and CONCOCT^35^ v1.1.0 and the resulting per-sample bin sets were then refined with DAS Tool^36^ v1.1.7. Per patient, bins were dereplicated with dRep^37^ v3.5.0, their completeness and contamination levels were determined with CheckM^38^ v1.0.2. Bin taxonomy was acquired with GTDB-Tk^39^ v2.5.2, and proteins were predicted with Prodigal^40^ v2.6.3.

### Read mapping and abundance determinations

To preclude the possibility of read mis-mapping, high-quality reads were first subset to 80 million and 15 million reads for the bulk metagenomic and VLP sequencing, respectively, and then mapped against the conglomerate of a patient’s contigs with Bowtie2^41–43^ v2.5.2. Then, to determine viral abundance, reads that mapped against viral contigs were subset and re-mapped to the species representatives. Similarly, bacterial reads were re-mapped to the final bin sets. These mappings were then used as input for three separate InStrain^44^ v1.8.0 analyses, one for phages in the VLP samples, another for phages in the bulk metagenomic samples, and a final one for bacterial bins.

### Inference of phage-host interactions

Phage-host interactions were inferred from Hi-C sequencing. Hi-C reads (average 115.3 million ± 113.8 million per sample, median 71.0 million) were adapter-trimmed with fastp^17–19^ v1.1.0 (--detect_adapter_for_pe) and mapped with bwa mem^46–48^ v0.7.19 (-5SP) to a combined reference composed of the phage contigs and the binned bacterial contigs. To prevent Hi-C reads mapping against prophages from being counted as concordant contacts, rather than discordant contacts expected from phage-host interactions, prophage regions were masked from the bacterial reference. The phage reference set was aligned against the binned bacterial contigs using BLAST^49^ v2.17.0+ and all hits longer than 5,000 bp were masked with BEDTools^50^ maskfasta v2.31.1. Mapped Hi-C reads were then normalised for contig length, depth, and distance to restriction sites using MetaCC^51^ v1.2.0 (-e Sau3AI -e MluCI --cover).

Phage-host interactions were inferred based on the distribution of Hi-C contacts between each contig and each bin, looking at two properties of Hi-C contacts: the contact breath, i.e. the number of contigs in a bacterial bin with which a contig had Hi-C contacts, and the contact intensity, i.e. the summed, normalized Hi-C contacts between a contig and all other contigs from a bin. A phage-host interaction had to have a contact breath of at least three and had to exceed a per-sample contact intensity cutoff. Intensity cutoffs were calibrated per sample on bacterial contigs, whose bin assignment is known independently. Each bacterial contig–bin pair was labelled positive if the contig belonged to that bin and negative otherwise, and after applying the breath criterion, the cutoff was set to the lowest intensity at which at least 95% of the pairs clearing it were true positives. Lastly, the distribution of Hi-C contacts along the viral contigs was manually screened, and phage-host interactions with unevenly distributed contacts were removed.

### Statistical analyses

All downstream statistical analyses and plotting were done using R v4.6.1 with the following packages: tidyverse, data.table, vegan^52^, patchwork, ggpubr, ggtext, ggrepel, colorspace, readxl, bipartite^53,54^, gridExtra, grid, paco^55^, ape, ggtree, glmmTMB^57,58^, DHARMa^59^, magick, scales, and pdftools. Unless otherwise specified, default settings were used for the command line tools above.

To test whether elevated nucleotide diversity in DGR-targeted genes was associated with a phage’s host range, log-transformed nucleotide diversity was modelled with a mixed-effects model in glmmTMB as a function of DGR targeting, the number of interacting hosts as a continuous predictor, expressed relative to a single host (n − 1), and the interaction between these. Nested random intercepts for coding sequence within phages and a random intercept for samples was added. A Student’s t response distribution with an identity link was used to accommodate the heavy-tailed diversity estimates, and the residual dispersion was allowed to differ between DGR-targeted and non-targeted genes.

Model adequacy was assessed using DHARMa. While the overall test of uniformity and the test of variance homogeneity for the number of interacting hosts predictor were significant, this was expected for a dataset of this size (7,313 observations from 2,113 coding sequences across 31 phages and 14 samples). Nvertheless, the quantile–quantile plot, the residual-versus-predictor plots, and the non-significant dispersion and outlier tests indicated no substantive violations, so the fit was judged acceptable. The significance of fixed effects was evaluated with Wald z-tests.

### AI declaration

Claude Opus 4.8 was used for copy editing and coding support, with edits and results verified and tested by authors.

## Results

### Hi-C sequencing enables high-confidence inference of phage-host interactions

As phage host ranges can shift over time, resolving them in the human gut requires following the same community across timepoints alongside culture-independent inference of phage–host interactions. We therefore drew on longitudinal faecal samples from our previously published faecal filtrate transplantation (FFT) randomised controlled clinical trial^16^. Briefly, participants received a laxative followed by either an FFT or a placebo consisting of a sterile saline infusion (**Fig. 1a**). Faecal samples were collected at 7 and 1 day prior to the intervention (days -7 and -1), and then on short intervals afterwards (days 2, 4, 7, 14, and 28). For this present study, we re-analysed the samples of two participants who received the placebo intervention, avoiding the effect of the FFT on the gut microbiome. Combining bulk metagenomic and VLP sequencing to capture bacteria, prophages, and free viral particles, we assembled a comprehensive view of the microbiome comprising 292 annotated *de novo* bacterial bins (>75% complete, <5% contamination). To focus on actively replicating viruses rather than cryptic prophages, we retained a viral species only if it was detected in the VLP fraction in one or more samples, yielding 365 high-quality *de novo* viral species (>90% complete, <5% contamination) across the two participants.

**Figure 1:**
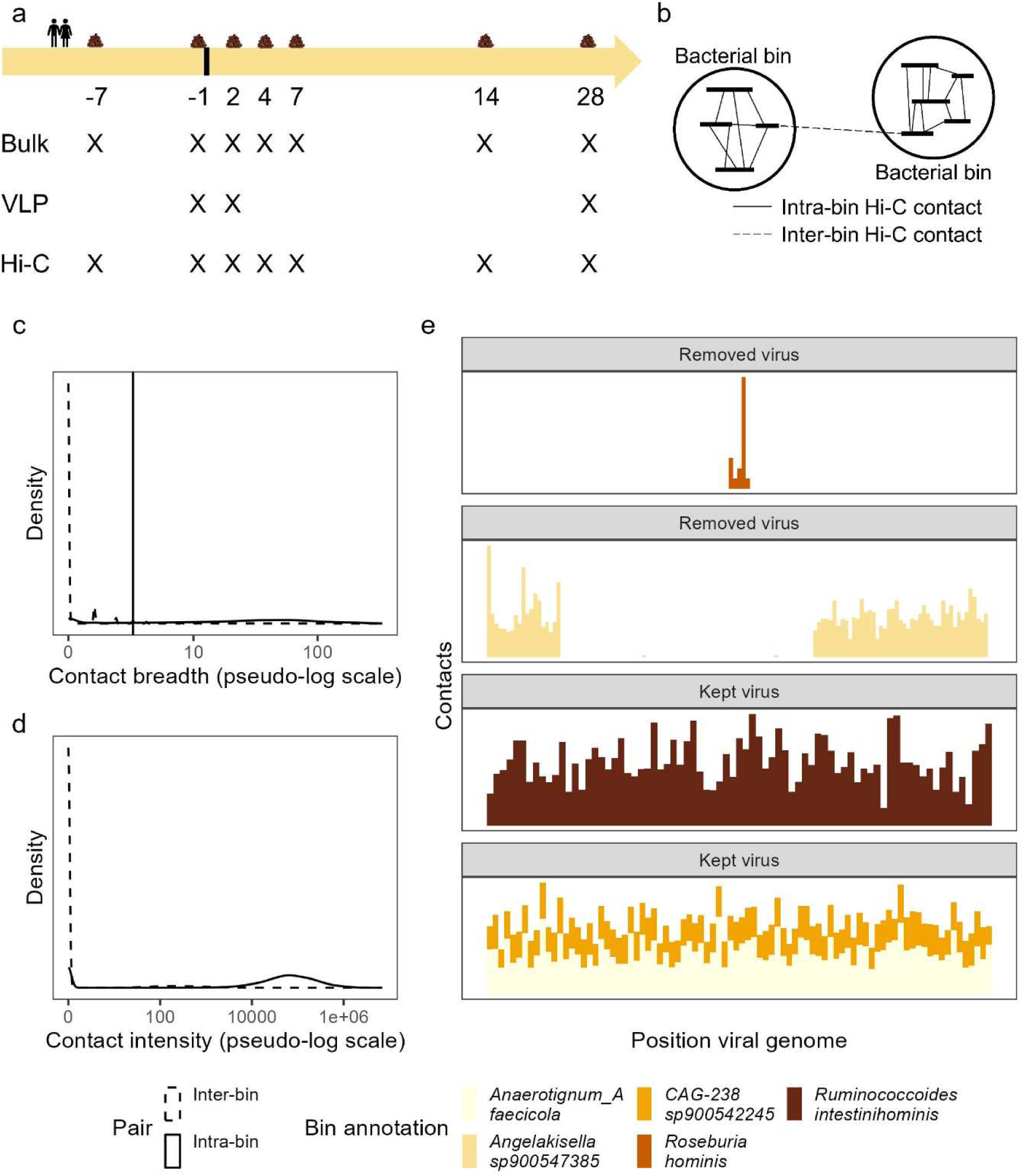
Sampling overview and validation of Hi-C contacts for inferring phage-host interactions. **a** Sampling scheme showing the timepoints at which bulk metagenomic, VLP and Hi-C sequencing was performed on fecal samples of the participants. **b** Schematic of expected Hi-C contacts: a contig is expected to share more contacts with contigs from its own bin (intra-bin) than with contigs from other bins (inter-bin). **c** Number of bacterial contigs in a bin with which a given contig shares Hi-C contacts (contact breath), for intra-bin and inter-bin contig–bin pairs (n = 13,90,460 pairs across all samples). **d** Distribution of summed, normalised Hi-C contacts (contact intensity) for intra-bin and inter-bin contig–bin pairs (n = 13,390,460 pairs across all samples). **e** Positions of Hi-C contacts along four example viral contigs: two viral contigs were removed because the Hi-C contacts were unevenly distributed along the viral genome, and two viral contigs that were retained.

We first assessed how the gut microbiome responded to the laxative, to establish whether these samples were suitable for tracking phage–host dynamics in the human gut. For both participants, bacterial richness decreased in the days immediately after the laxative (days 2, 4, and 7), before returning to baseline by day 14 (**Supp. Fig. 1a**). Bacterial community composition differed between the days immediately after the laxative and the remaining timepoints (days −7, −1, 14, and 28), explaining 6% of the variation (**Supp. Fig. 1b**; PERMANOVA of Bray-Curtis dissimilarities stratified by participant, 999 permutations, R² = 0.06, F = 1.68, p = 0.005), while inter-participant differences explained the majority of the variation (R² = 0.57). Bulk-derived viral richness tracked bacterial richness (two-sided Spearman’s ρ = 0.91, p < 0.001) and showed the same transient decrease, whereas VLP-derived viral richness did not show a significant association (two-sided Spearman’s ρ = −0.60, p = 0.24), consistent with the transient loss of bacterial species and their integrated prophages, but not of free viral particles (**Supp. Fig 1c**). The bulk viral community composition as a whole did not shift significantly between the early post-laxative days and the other timepoints (**Supp. Fig. 1d**; PERMANOVA of Bray-Curtis dissimilarities stratified by participant, 999 permutations, R² = 0.05, F = 1.05, p = 0.42), while inter-participant differences some variation (R² = 0.39). Taken together, the laxative caused only a transient, minor perturbation of the gut bacteria with no detectable effect on the viral community, making this dataset suitable for tracking the dynamics of phage–host interactions without the confounding effect of a major, sustained disturbance of the gut microbiome.

Phage–host interactions were inferred using Hi-C sequencing. We reasoned that an infecting phage would interact similarly with a bacterial bin as the bacterial contigs belonging to that bin. As the bin membership of bacterial contigs is known, contig–bin pairs in which the contig belongs to the bin (intra-bin) served as true interactions, while pairs in which the contig did not (inter-bin) as false interactions (**Fig. 1b**). First, we examined the distribution of Hi-C contacts between intra- and inter-bin pairs, looking at two properties: contact breath, i.e. the number of contigs in a bin with which a contig had Hi-C contacts, and contact intensity, i.e. the summed, normalized Hi-C contacts between a contig and all other contigs from a bin. Inter-bin pairs had a median contact breath of 0 (IQR 0) and contact intensity of 0 (IQR = 0), while intra-bin pairs had a median contact breath of 27 (IQR = 54) and a contact intensity of 50,002 (IQR = 120,674) (**Fig. 1cd**).

We used this separation to set cutoffs for calling a contig–bin interactions. First, we required a contact breadth of at least three, which removed 96.2% of inter-bin pairs and 21.1% of intra-bin pairs. Second, an interaction also needed to exceed a per-sample contact intensity cutoff, which was calibrated per sample to achieve a precision of at least 95%, meaning that 95% of called interactions were intra-bin pairs (**Supp. Fig 2**). Together, these cutoffs resulted an overall recall of 77.0% pooled across samples, meaning that 77% of intra-bin pairs were recovered. Both these cutoffs were then applied to viral contigs and bacterial bins, yielding 483 phage-host interactions between 106 phages and 99 bacterial bins. Lastly, we also did an additional, manual screen of where these contacts mapped on the viral contigs. We reasoned that for a genuine interaction, contacts should be evenly distributed along the viral contig, rather than confined to a specific region, which could indicate mismapping or assembly errors (**Fig. 1e**). This screen excluded 9 phages, resulting in high-confidence 380 phage-host interactions between 97 phages and 76 bacterial bins.

### Phage-host interaction networks are modular and congruent

Overall, we identified a host for 97 of 365 (26.6%) phages. Most phages interacted with a single bacterial species, however 8 of 97 (8.2%) phages interacted with multiple bacterial species, up to a maximum of four (**Fig. 2a**). As samples were collected longitudinally, we were able to follow phage-host interactions over time (**Fig. 2b**). Some interactions were detected at every timepoint, such as between an unclassified *Caudoviricetes* phage and *Acidaminococcus intestine*. Other phages interacted more sporadically with their hosts, such as *Endlipuvirus* interacting with *Alistipes putredinis* or *Carjivirus* interacting with the known *Crassvirales* host *Phocaeicola vulgatus*^60^. There were also phages that interacted with multiple hosts, such as another unannotated *Caudoviricetes*, which was found to interact with both *Mediterraneibacter torques* and *Enterocloster sp000431375,* both *Lachnospiraceae*.

**Figure 2:**
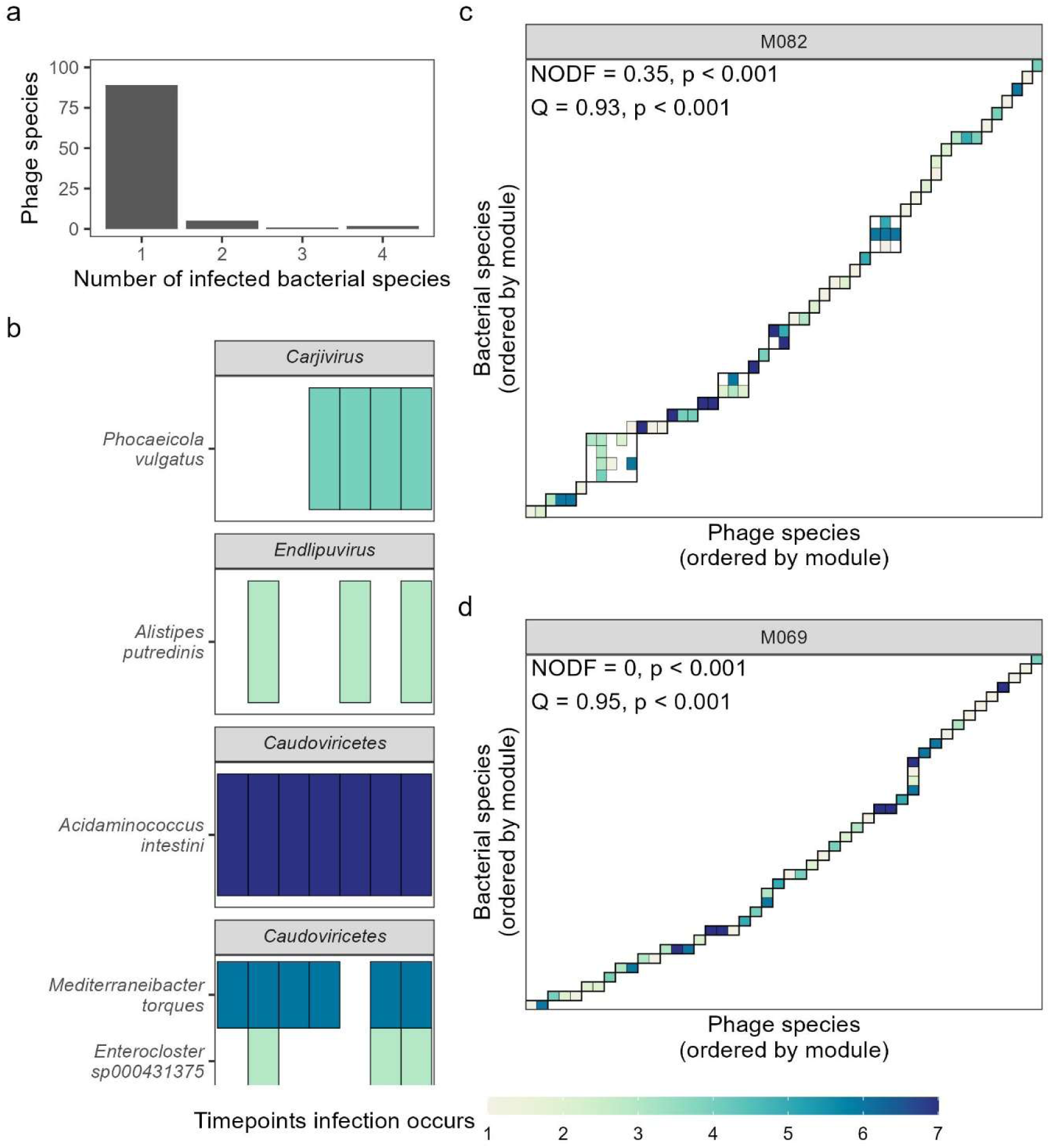
Phage-host interaction networks. **a** Number of host species each phage species interacted with (n = 97 phage species). **b** Phage-host interactions over time of four example. Each tile indicates that an interaction was detected at that timepoint, with the colour indicating the total number of timepoints that phage-host interaction was detected. **c.** Phage-host interaction network of participant M082. Grey boxes indicate the modules. NODF = nestedness; Q = modularity. **d.** As in c, for participant M069.

To characterise the overall structure of these interactions, we represented them as a bipartite network for each participant, in which phages and bacteria form two distinct sets of nodes and edges connect each phage to the bacteria it interacts with (**Fig. 2c, d**). These networks are often described as nested or modular^61^. Nested networks are characterized by a hierarchical structure in which broad host-range phages infect many bacteria, including those infected by only a few phages, while narrow host-range phages infect only the bacteria that are infected by many phages (**Supp. Fig. 3a**). In modular networks, phages and their hosts form distinct groups or modules, with many interactions within the modules and few between modules (**Supp. Fig. 3b**). Both networks were significantly anti-nested (999 permutations with quantitative null-model with fixed row and column totals, p < 0.001 for each participant). This may reflect the significant modularity of the networks (Dormann & Strauss, 999 permutations with quantitative null-model with fixed row and column totals, p < 0.001 for each participant). The network of participant M082 partitioned into several modules (**Fig. 2c and Supp. Fig. 3b**), whereas the network of participant M069 consisted of many small, primarily single-phage-single-host pairs, which are maximally modular by definition (**Fig. 2d**).

As phages are dependent on their bacterial hosts, their phylogeny often mirrors that of theirs hosts, which is called congruency^62^. We examined whether the detected phage-host interactions followed this pattern using a Procrustes Application to Cophylogenetic Analysis (PACo) based on their taxonomies. For both participants, phage-host interactions were significantly congruent (PACo, 999 permutations with quantitative null-model with fixed row totals, separated by participant, p < 0.001), consistent with co-evolution between phages and their bacterial hosts.

### Human gut phages encode DGRs

Of the 97 phages for which we identified a bacterial host, 31 (32.0%) encoded a DGR (**Fig. 3a**). DGRs have been divided into six different clades based on the phylogeny of their reverse transcriptase^13^. Consistent with what was previously observed in human gut viruses, most of these phages encoded a DGR from clades 1 and 4^13^, with one phage encoding a clade 3 DGR, (**Fig. 3a, Supp. Fig. 4**). The majority of DGRs targeted one gene, although some also targeted an additional one, with these genes predicted to encode tail (fibre) and Hoc-like head decoration proteins, with different clades preferentially targeting different genes, also as previously observed^13^ (**Fig. 3b**; two-sided Fisher’s exact test, corrected Cramér’s V = 0.49, p = 0.006). Upon further examination, however, the Hoc-like head decoration proteins resembled the recently described tentaclins proteins^63^. Tentaclins were defined as having a VR towards their 5’-end and a N-terminal DUF6273 domain, which is similar to C-type lectin (C-lec) domain, followed by one or more Ig-like domains^63^. Similarly, the Hoc-like head decoration genes found in this study also encoded their VR more towards the the 5′ rather than the 3′ end of the gene and all (17/17) encoded N-terminal DUF6273 domains, for two of which we were also able to detect Ig-like domains (**Supp. Fig 5**). Therefore, we decided to reclassify these as putative tentaclins. While the exact function of tentaclins is not yet known, they are speculated to act as receptor-binding proteins, as well.

**Figure 3:**
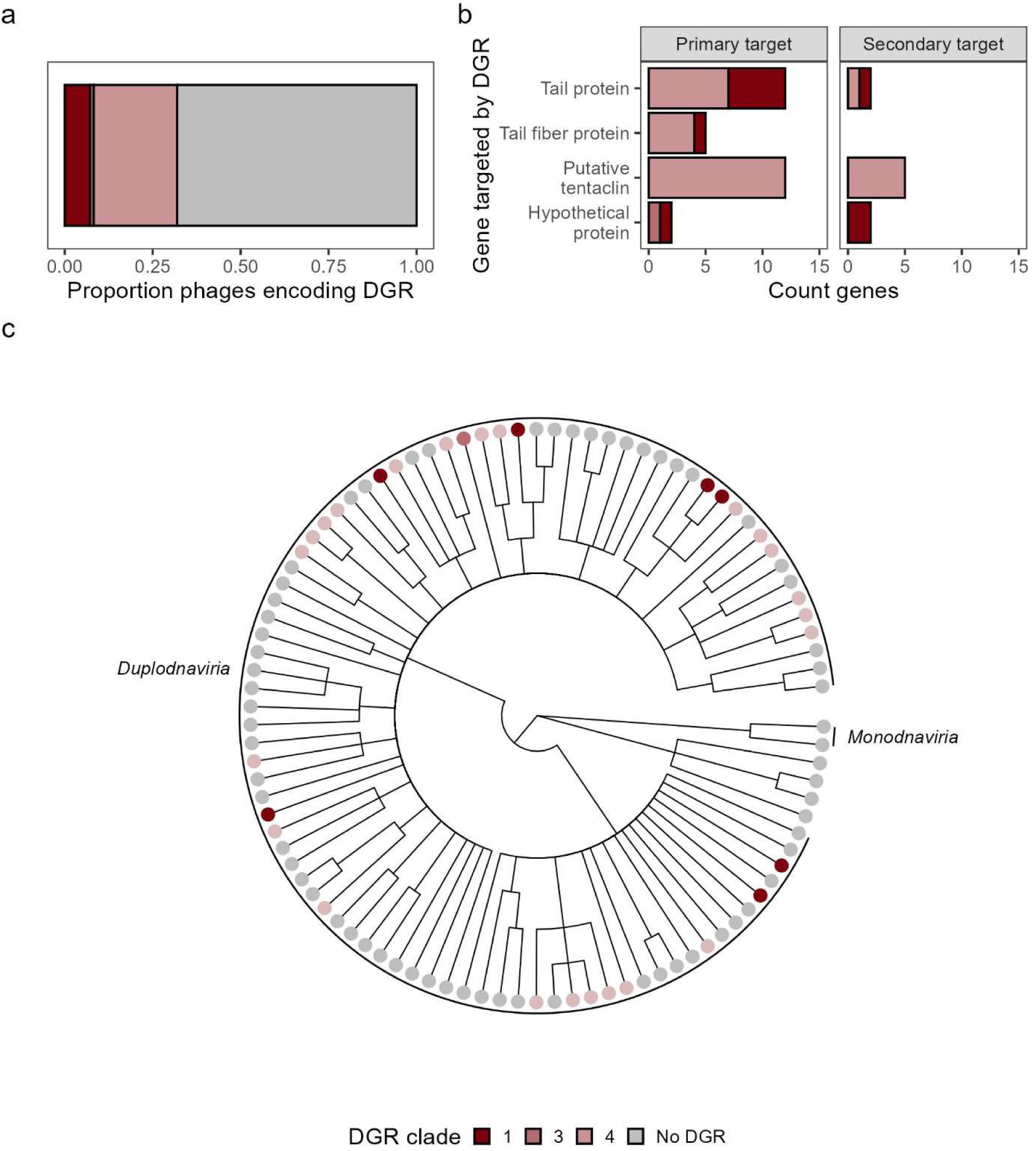
DGRs in human gut viruses. **a.** Proportion of phages that encode DGRs, coloured by DGR clade (n = 97 phages). **b.** Gene annotations of primary DGR targets, coloured by DGR clade and separated by whether they are the primary or secondary DGR target (n = 31 phages). **c.** Dendrogram of gut phages, coloured by encoded DGR clade (n = 40 DGR-targeted genes).

DGRs were encoded by diverse taxa within *Duplodnaviria*, and DGR clades were not confined to any particular viral lineage, either (**Fig. 3c**). As DGR reverse transcriptases are monophyletic^13^, this suggests that DGRs may have disseminated through horizontal gene transfer.

### Hypermutation of target genes by DGRs correlates with phage host ranges

As we hypothesized that DGRs enable phages to expand their host range, we examined whether DGR-encoding phages interacted with more bacterial species than those that did not encode a DGR. Phages encoding DGRs were more likely to interact with multiple bacterial species compared to those that did not encode a DGR (**Fig. 4a**; two-sided Fisher’s exact test, odds ratio = 18.4, p = 0.001). To test whether this related to the activity of DGRs, we modelled the nucleotide diversity of phage genes as a function of DGR targeting, the number of hosts a phage interacted with at each timepoint, and the interaction between these, in a mixed-effects model (**Fig 4b**). When interacting with a single host, DGR-targeted genes had a 47% higher nucleotide diversity than non-DGR targeted genes (two-sided Wald z-test, β = 0.38, SE = 0.05, z = 7.69, p < 0.001). Each additional host was associated with a further 22.5% increase in the diversity of DGR-targeted genes (β = 0.20, 95% CI 0.05% – 0.36%), whereas nucleotide diversity in non-DGR targeted genes did not appear to be affected (β = -0.004, 95% CI -0.02% – 0.02%), with the two slopes differing significantly (two-sided Wald z-test, β = 0.21, SE = 0.08, z = 2.57, p = 0.01). Therefore, multi-host interactions were associated with elevated diversity in the DGR-targeted genes, specifically, rather than across the phage genome in general.

**Figure 4:**
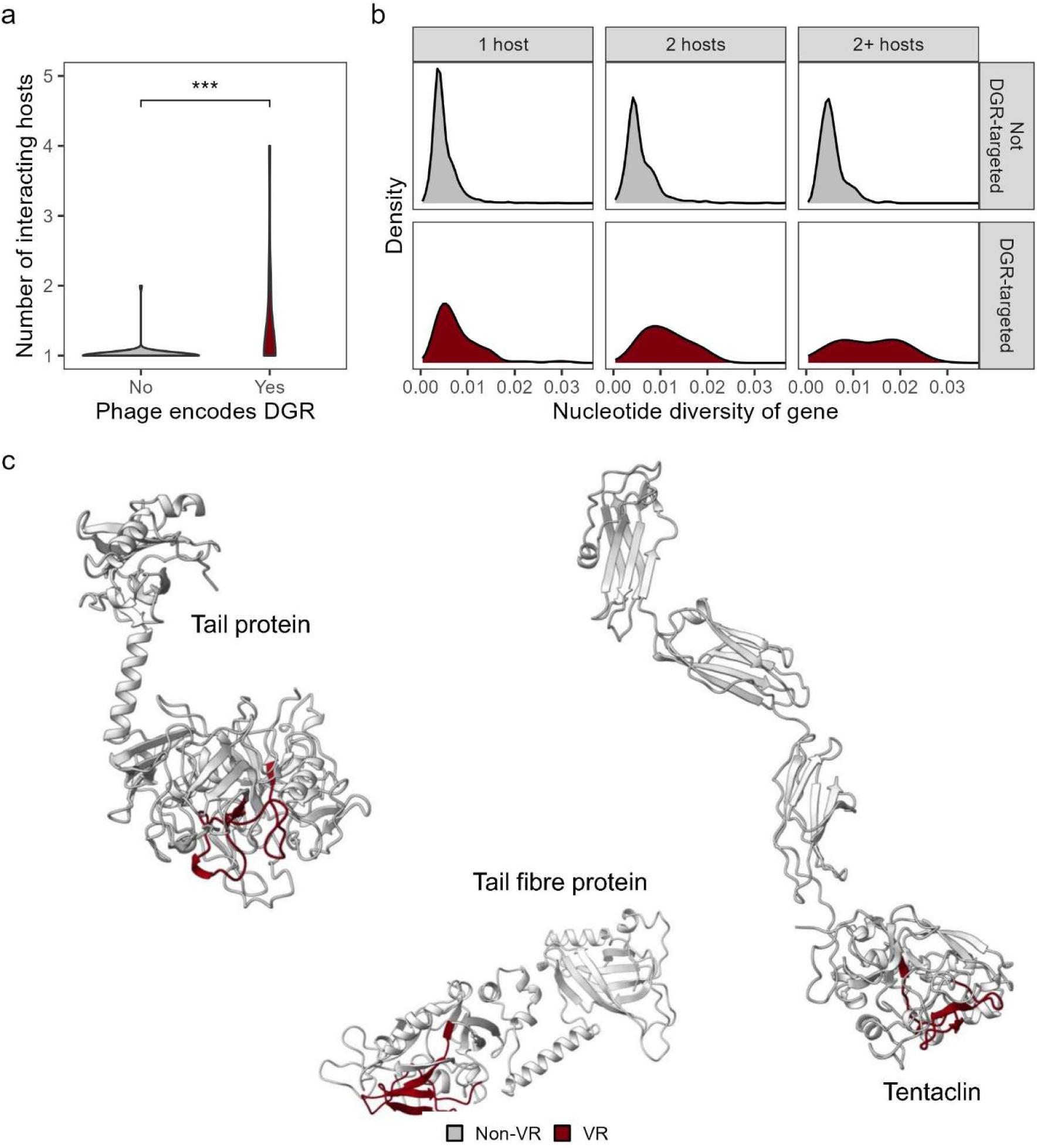
Host range of DGR-encoding phages. **a.** Number of interacting bacterial species per phage, stratified by whether the phage encodes a DGR (n = 97 phages). **b.** Nucleotide diversity of genes of DGR-encoding phages, stratified by whether the gene is targeted by a DGR and the number of interacting bacterial species of that phage per timepoint (n = 7,313 genes with nucleotide diversities > 0 over all timepoints, from 2,113 unique genes). **c.** 3D protein models of three DGR-targeted genes. The region targeted by the DGR, the VR, is coloured (n = 3 proteins).

As expected, given that VRs are by definition the target of DGR-hypermutation, mutations were heavily concentrated here with 3,179 of 3,910 (81.3%) non-synonymous mutations falling within the VRs, despite these encompassing only 9,081 of 116,277 (7.8%) bp of the DGR-targeted gene sequence, a 51-fold enrichment in non-synonymous mutations compared to the rest of the gene. To affect host recognition, we reasoned that these hypermutated VRs should be located on the surface-exposed region of the protein in order to interact with the host. We therefore predicted the protein structures for a tail, a tail fibre, and a putative tentaclin protein. Indeed, in each case, the VR was located on the exposed protein surface (**Fig. 4c**). Taken together, these results show that DGRs hypermutate the surface-exposed regions of phage receptor-binding proteins and that this DGR-driven diversity scales with the number of hosts a phage interacts with.

## Discussion

Phage host ranges are increasingly recognized as being broader and more dynamic than one phage infecting one single host^5–8^. DGRs may be one of the mechanisms through which phages achieve this, however, direct evidence has been limited. By combining longitudinal sampling with bulk metagenomic, VLP, and Hi-C sequencing, our study resolves phage-host interactions directly in the human gut, addressing this gap.

The overall structure of phage-host interactions was both modular and taxonomically congruent: phages tended to interact with hosts within their own module, and phage taxonomy broadly tracked host taxonomy. This is the expected pattern if phages and their hosts co-evolve, whereby phages, as parasites dependent on their bacterial hosts, evolve in tandem with them^62^. Phage-host infection networks are often reported as nested rather than modular, but this is primarily seen in networks of closely related phages and hosts, i.e. primarily in networks including multiple strains of the same phage and bacterial species^64^. By contrast, networks spanning multiple phage and bacterial species, as in our study, more often show a modular structure^65–67^. Such modularity is thought to arise from co-evolution together with the general specificity of phage infection^65^. This was also observed here, with most phages interacting with a single bacterial species, and those that interacted with multiple species preferentially did so with closely related taxa.

DGR-encoding phages were abundant in our samples, with around a third of phages encoding a DGR. These DGRs were encoded by various taxa across *Duplodnaviria* and did not cluster within specific viral lineages. This lack of clustering, combined with the previously reported monophyly of the reverse transcriptase of DGRs^13^, suggests that DGRs may have spread through horizontal gene transfer. Concordant with the paradigm of phage specificity, we mostly found phages linked to a single bacterial species, however those encoding a DGR were more likely to interact with multiple species. Mechanistically, DGR-mediated host-range expansion would occur through mutations in the VR of the target genes, usually receptor-binding proteins^12^. Accordingly, DGR-targeted genes in our study showed higher nucleotide diversity and, importantly, this effect depended on the host range of the phage, being more pronounced in phages interacting with multiple bacterial hosts. Non-synonymous mutations were concentrated in the VR, which was located on the surface-exposed region in each predicted protein structure, where mutations would alter the binding specificity or affinity without destabilising its overall structure.

Notably, although multi-host phages were more likely to encode a DGR, most DGR-encoding phages interacted with a single bacterial species during our study. We propose two possibilities for this. First, given that we inferred phage-host interactions based on the assumption that infecting phages would interact similarly with the bacterial bins as the bacterial contigs belonging to that bin, we would have missed phage-host interactions that were weaker or more transient than those occurring within bacterial bins. Second, phages also use DGRs to adapt to variation within a single bacterial species, such as when infecting multiple strains of the same species or when adapting to phenotypic switches of their host, as is the case in the canonical *Bordetella* phage^12^. Our study could not capture either process, as bacterial genomes were resolved at the species level (95% ANI), making strain-level host switching invisible, and phenotypic switching cannot be detected by sequencing at all.

A limitation of our study is that the sampling was performed on two participants only. However, to resolve phage-host interactions over time and associate these with the diversity of DGR-target genes, the combination of deep bulk metagenomic, VLP, and Hi-C sequencing was necessary, which would have been unfeasible in a larger-scale cohort. Our approach nevertheless recovered a sufficient number of phages, allowing us to associate their diversity in DGR-targeted genes with their host range through mixed-effect models, while accounting for the repeated sampling of the phages. Our results should therefore be interpreted as properties of these phages, with larger cohorts being necessary to generalize these findings across gut viromes.

A second limitation is that our study does not capture phage-host infections per se, but rather physical interactions between the phage and host DNA. Nevertheless, co-localization of phage and host DNA reflects the entry of the phage genome into the host cell, which is the essential first step of any successful phage infection. Furthermore, all phages included in our study were present in the VLP fraction, and while this did not discern from which host these originated, it does confirm that these phages are actively replicating.

In conclusion, by combining longitudinal sampling with culture-independent inference of phage-host interactions, we find *in situ* evidence that DGR-driven hypermutation of the surface-exposed regions of phage receptor-binding proteins is associated with phage host range. This highlights DGRs as a candidate mechanism for host-range adaptation and expansion, of interest for applications such phage therapy.

## Acknowledgements

We would like to thank Ivan Liachko and Ben Auch (Phase Genomics Inc, Seattle, Washington, USA) for their assistance and advice with Hi-C sequencing. We are grateful to Simon Roux (Joint Genome Institute, Berkeley, California, USA) for his excellent correspondence and help with the clade assignment of the DGRs and Julie Boulbes (Institut Pasteur, Paris, France) for her suggestions on host assignments based on Hi-C data.

A.W.M.vdH is supported by an Amsterdam UMC PhD Scholarship (2022). M.Z. is supported by HORIZON-MSCA-2025-PF (2025, 101276727). M.N. is supported by a personal ZonMw Vici grant (2020; 09150182010020) and the European Research Council Advanced Grant FATGAP (101141346); P.A.dJ. is supported by a personal Veni grant (2024; 232290).

---

**Supplementary Figure 1:**
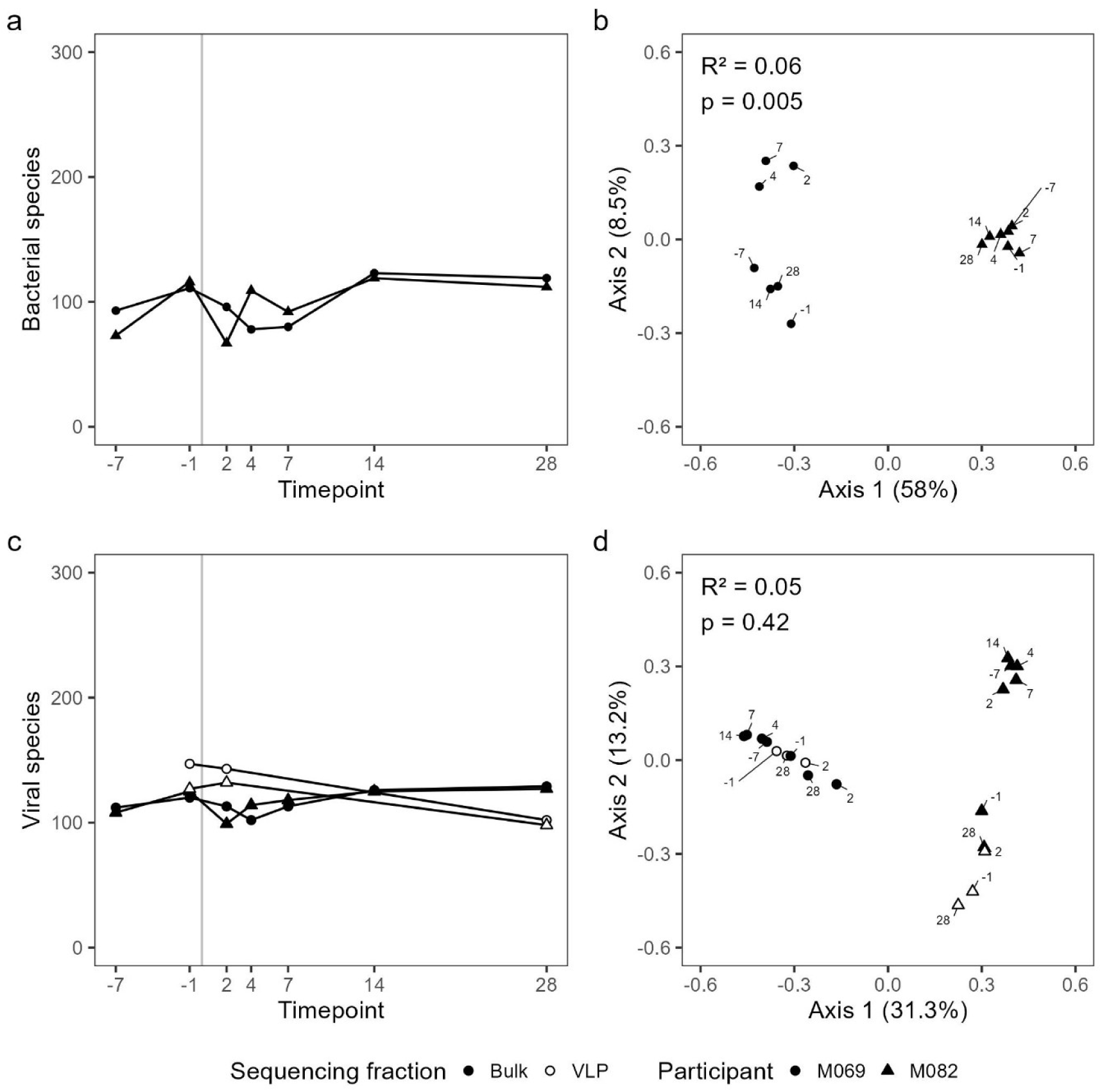
Bacterial and viral community dynamics in two participants over time. **a** Bacterial species richness at each timepoint. **b** Principal coordinates analysis (PCoA) of bacterial community composition based on Bray-Curtis dissimilarities. **c** Viral species richness at each timepoint in both the bulk metagenomes and viral-like particle (VLP) viromes. **d** PCoA of viral community composition in bulk metagenomes and VLP viromes based on Bray–Curtis dissimilarity. R^2^ and p-values refer to the bulk-derived viral community, no statistical test was run on the VLP-derived viral communities (n = 14 bulk metagenomes and n = 6 VLP viromes, across the two participants).

**Supplementary Figure 2:**
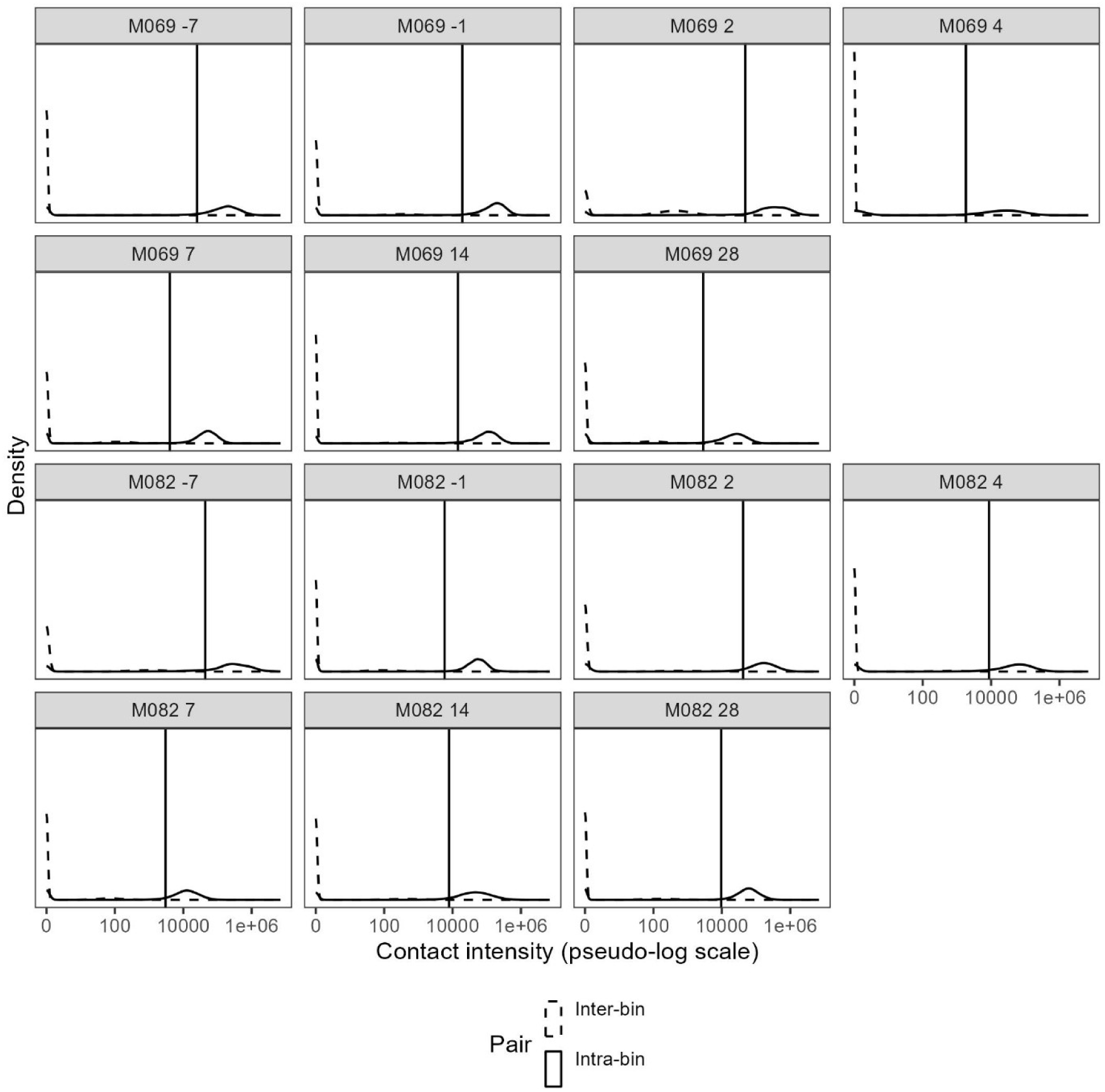
Distribution of Hi-C contacts and cutoffs. Per-sample distribution of summed, normalised Hi-C contacts (contact intensity) for intra-bin and inter-bin contig–bin pairs. Per-sample cutoff of the contact intensity is indicated with a vertical line. (M069 -7 n = 795,615; M069 -1 n = 1,176,156; M069 2 n = 818,592; M069 4 n = 626,262; M069 7 n = 607,680; M069 14 n = 1,383,750; M069 28 n = 1,227,247; M082 -7 n = 514,212; M082 -1 n = 1,312,424; M082 2 n = 376,339; M082 4 n = 1,168,698; M082 7 n = 829,472; M082 14 n = 1 320,781; M082 28 n = 1,233,232)

**Supplementary Figure 3:**
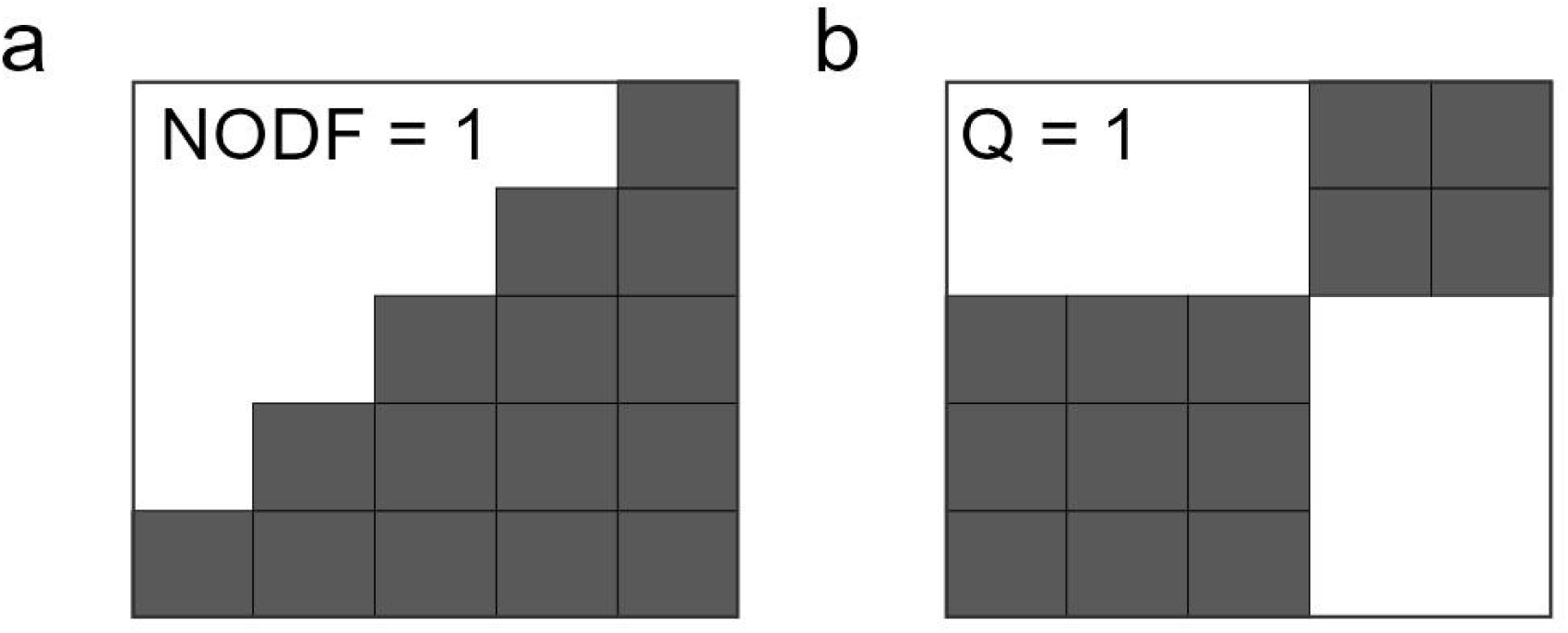
Examples of bipartite network structures. **a** Perfectly nested bipartite network. **b** Perfectly modular bipartite network.

**Supplementary Figure 4:**
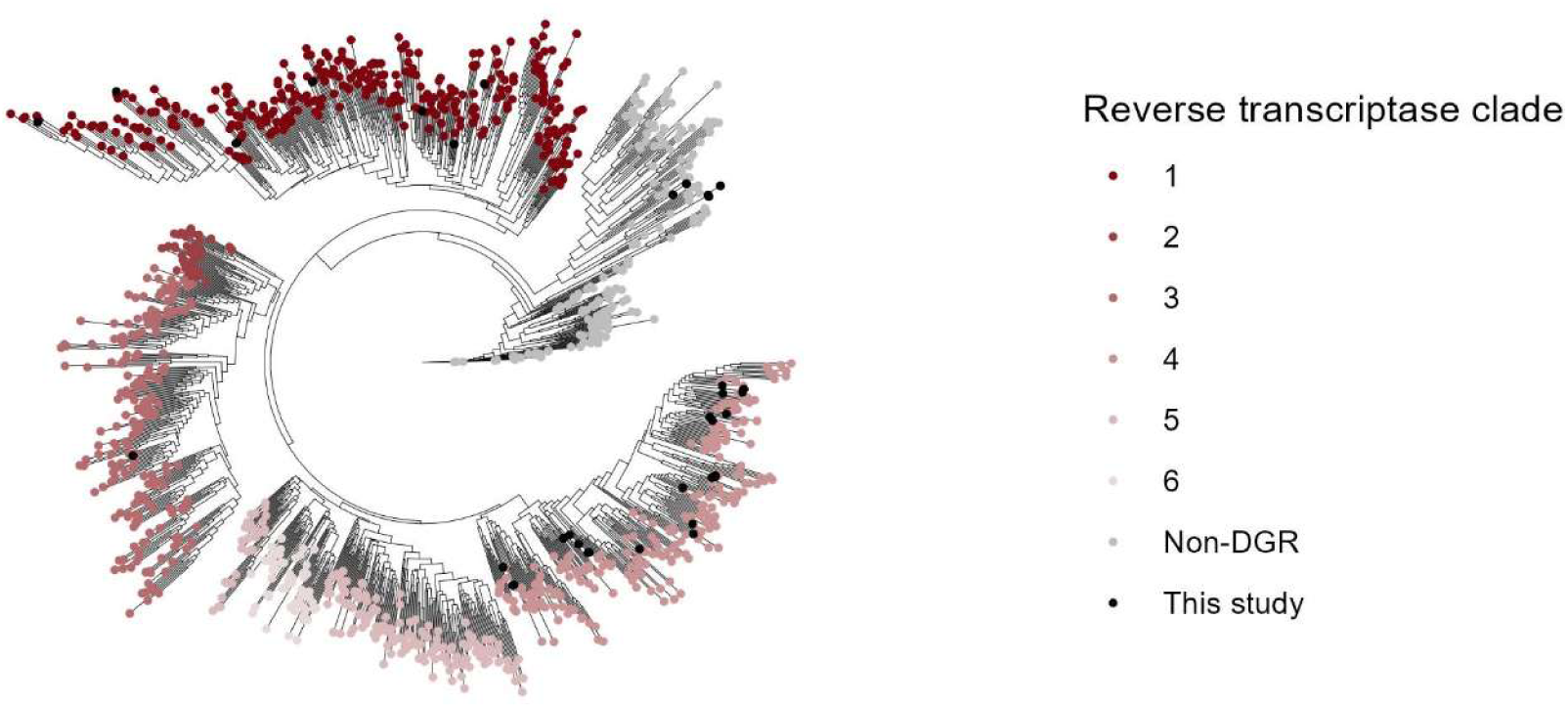
Phylogenetic tree of reverse transcriptase. A phylogenetic tree of the reverse transcriptase from the study by Roux et al., with the reverse transcriptase within phages linked to a bacterial host from this study placed on the tree in black. DGR clades were taken from Roux *et al*. and the clades of the reverse transcriptase found in this study were annotated based on where they clustered (n = 1,250 reverse transcriptases).

**Supplementary Figure 5:**
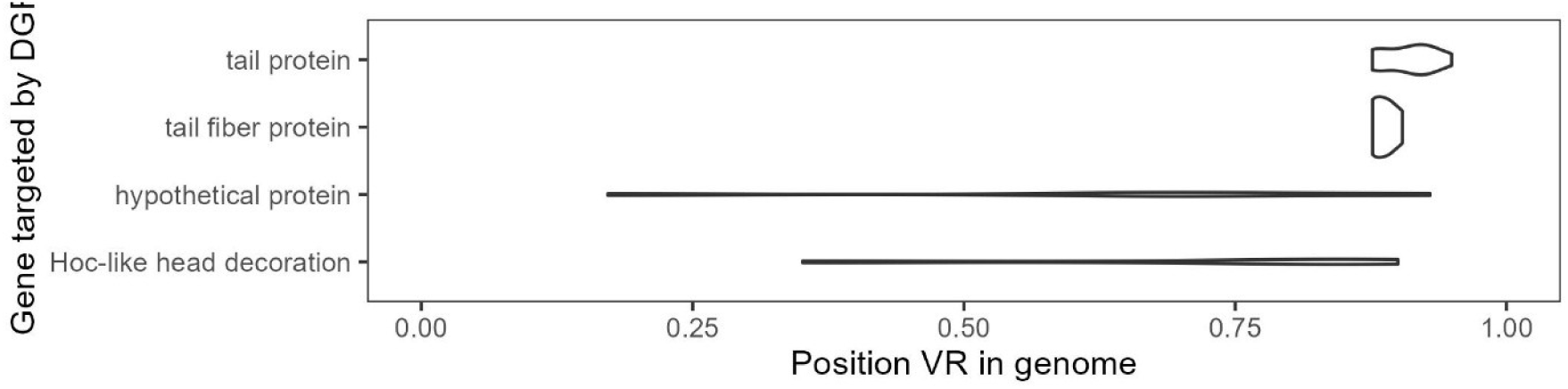
Position of VR in DGR-targeted genes. Separated by gene annotation. Hoc-like head decoration genes were reclassified as putative tentaclins (n = 40 DGR-targeted genes).

## References

1. Chevallereau, A., Pons, B. J., van Houte, S. & Westra, E. R. Interactions between bacterial and phage communities in natural environments. Nat Rev Microbiol 20, 49–62 (2022).

2. Shkoporov, A. N., Turkington, C. J. & Hill, C. Mutualistic interplay between bacteriophages and bacteria in the human gut. Nat Rev Microbiol 20, 737–749 (2022).

3. Gordillo Altamirano, F. L. & Barr, J. J. Phage Therapy in the Postantibiotic Era. Clinical Microbiology Reviews 32, 10.1128/cmr.00066-18 (2019).

4. Xu, Y. Phage and phage lysins: New era of bio-preservatives and food safety agents. Journal of Food Science 86, 3349–3373 (2021).

5. Hwang, Y., Roux, S., Coclet, C., Krause, S. J. E. & Girguis, P. R. Viruses interact with hosts that span distantly related microbial domains in dense hydrothermal mats. Nat Microbiol 8, 946–957 (2023).

6. Bignaud, A. et al. Phages with a broad host range are common across ecosystems. Nat Microbiol 10, 2537–2549 (2025).

7. de Jonge, P. A., Nobrega, F. L., Brouns, S. J. J. & Dutilh, B. E. Molecular and Evolutionary Determinants of Bacteriophage Host Range. Trends Microbiol 27, 51–63 (2019).

8. Leprince, A., Somerville, V., Addablah, A. A., Morency, C. & Moineau, S. Phage host range: determinants, dynamics and applications. Nat Rev Microbiol 1–21 (2026) doi:10.1038/s41579-026-01306-x.

9. Plattner, M. et al. Structure and Function of the Branched Receptor-Binding Complex of Bacteriophage CBA120. J Mol Biol 431, 3718–3739 (2019).

10. Schwarzer, D. et al. A multivalent adsorption apparatus explains the broad host range of phage phi92: a comprehensive genomic and structural analysis. J Virol 86, 10384–10398 (2012).

11. Tu, J. et al. Dual host specificity of phage SP6 is facilitated by tailspike rotation. Virology 507, 206–215 (2017).

12. Liu, M. et al. Reverse Transcriptase-Mediated Tropism Switching in Bordetella Bacteriophage. Science 295, 2091–2094 (2002).

13. Roux, S. et al. Ecology and molecular targets of hypermutation in the global microbiome. Nat Commun 12, 3076 (2021).

14. Camargo, A. P. et al. A genomic atlas of the human gut virome elucidates genetic factors shaping host interactions. 2025.11.01.686033 Preprint at 10.1101/2025.11.01.686033 (2025).

15. Benler, S. et al. A diversity-generating retroelement encoded by a globally ubiquitous Bacteroides phage. Microbiome 6, 191 (2018).

16. Wortelboer, K. et al. Phage-microbe dynamics after sterile faecal filtrate transplantation in individuals with metabolic syndrome: a double-blind, randomised, placebo-controlled clinical trial assessing efficacy and safety. Nat Commun 14, 5600 (2023).

17. Chen, S. Ultrafast one-pass FASTQ data preprocessing, quality control, and deduplication using fastp. iMeta 2, e107 (2023).

18. Chen, S. fastp 1.0: An ultra-fast all-round tool for FASTQ data quality control and preprocessing. iMeta 4, e70078 (2025).

19. Chen, S., Zhou, Y., Chen, Y. & Gu, J. fastp: an ultra-fast all-in-one FASTQ preprocessor. Bioinformatics 34, i884–i890 (2018).

20. Bushnell, B. BBMap - Official BBTools Suite. https://bbmap.org/.

21. Li, D. et al. MEGAHIT v1.0: A fast and scalable metagenome assembler driven by advanced methodologies and community practices. Methods 102, 3–11 (2016).

22. Li, D., Liu, C.-M., Luo, R., Sadakane, K. & Lam, T.-W. MEGAHIT: an ultra-fast single-node solution for large and complex metagenomics assembly via succinct de Bruijn graph. Bioinformatics 31, 1674–1676 (2015).

23. Guo, J. et al. VirSorter2: a multi-classifier, expert-guided approach to detect diverse DNA and RNA viruses. Microbiome 9, 37 (2021).

24. Shen, W., Le, S., Li, Y. & Hu, F. SeqKit: A Cross-Platform and Ultrafast Toolkit for FASTA/Q File Manipulation. PLoS One 11, e0163962 (2016).

25. Nayfach, S. et al. CheckV assesses the quality and completeness of metagenome-assembled viral genomes. Nat Biotechnol 39, 578–585 (2021).

26. Zielezinski, A. et al. Ultrafast and accurate sequence alignment and clustering of viral genomes. Nat Methods 22, 1191–1194 (2025).

27. Bouras, G. et al. Pharokka: a fast scalable bacteriophage annotation tool. Bioinformatics 39, btac776 (2023).

28. Protein structure-informed bacteriophage genome annotation with Phold | Nucleic Acids Research | Oxford Academic. https://academic.oup.com/nar/article/54/1/gkaf1448/8415830?login=false.

29. Eddy, S. R. Accelerated Profile HMM Searches. PLOS Computational Biology 7, e1002195 (2011).

30. Katoh, K., Misawa, K., Kuma, K. & Miyata, T. MAFFT: a novel method for rapid multiple sequence alignment based on fast Fourier transform. Nucleic Acids Res 30, 3059–3066 (2002).

31. Kozlov, A. M., Darriba, D., Flouri, T., Morel, B. & Stamatakis, A. RAxML-NG: a fast, scalable and user-friendly tool for maximum likelihood phylogenetic inference. Bioinformatics 35, 4453–4455 (2019).

32. Barbera, P. et al. EPA-ng: Massively Parallel Evolutionary Placement of Genetic Sequences. Syst Biol 68, 365–369 (2019).

33. Czech, L., Barbera, P. & Stamatakis, A. Genesis and Gappa: processing, analyzing and visualizing phylogenetic (placement) data. Bioinformatics 36, 3263–3265 (2020).

34. Kang, D. D. et al. MetaBAT 2: an adaptive binning algorithm for robust and efficient genome reconstruction from metagenome assemblies. PeerJ 7, e7359 (2019).

35. Alneberg, J. et al. Binning metagenomic contigs by coverage and composition. Nat Methods 11, 1144–1146 (2014).

36. Sieber, C. M. K. et al. Recovery of genomes from metagenomes via a dereplication, aggregation and scoring strategy. Nat Microbiol 3, 836–843 (2018).

37. Olm, M. R., Brown, C. T., Brooks, B. & Banfield, J. F. dRep: a tool for fast and accurate genomic comparisons that enables improved genome recovery from metagenomes through de-replication. ISME J 11, 2864–2868 (2017).

38. Parks, D. H., Imelfort, M., Skennerton, C. T., Hugenholtz, P. & Tyson, G. W. CheckM: assessing the quality of microbial genomes recovered from isolates, single cells, and metagenomes. Genome Res 25, 1043–1055 (2015).

39. GTDB-Tk: a toolkit to classify genomes with the Genome Taxonomy Database | Bioinformatics | Oxford Academic. https://academic.oup.com/bioinformatics/article/36/6/1925/5626182.

40. Hyatt, D. et al. Prodigal: prokaryotic gene recognition and translation initiation site identification. BMC Bioinformatics 11, 119 (2010).

41. Langmead, B. & Salzberg, S. L. Fast gapped-read alignment with Bowtie 2. Nat Methods 9, 357–359 (2012).

42. Langmead, B., Trapnell, C., Pop, M. & Salzberg, S. L. Ultrafast and memory-efficient alignment of short DNA sequences to the human genome. Genome Biol 10, R25 (2009).

43. Langmead, B., Wilks, C., Antonescu, V. & Charles, R. Scaling read aligners to hundreds of threads on general-purpose processors. Bioinformatics 35, 421–432 (2019).

44. Olm, M. R. et al. inStrain profiles population microdiversity from metagenomic data and sensitively detects shared microbial strains. Nat Biotechnol 39, 727–736 (2021).

45. Zhu, K. et al. Proof-of-concept of host attribution of antimicrobial resistance genes using wastewater Hi-C metagenome sequencing. J Water Health 24, 148–159 (2025).

46. Li, H. Aligning sequence reads, clone sequences and assembly contigs with BWA-MEM. Preprint at 10.48550/arXiv.1303.3997 (2013).

47. Li, H. & Durbin, R. Fast and accurate short read alignment with Burrows-Wheeler transform. Bioinformatics 25, 1754–1760 (2009).

48. Li, H. & Durbin, R. Fast and accurate long-read alignment with Burrows-Wheeler transform. Bioinformatics 26, 589–595 (2010).

49. Camacho, C. et al. BLAST+: architecture and applications. BMC Bioinformatics 10, 421 (2009).

50. Quinlan, A. R. & Hall, I. M. BEDTools: a flexible suite of utilities for comparing genomic features. Bioinformatics 26, 841–842 (2010).

51. Du, Y. & Sun, F. MetaCC allows scalable and integrative analyses of both long-read and short-read metagenomic Hi-C data. Nat Commun 14, 6231 (2023).

52. Oksanen, J., et al. Vegan: Community Ecology Package. (2025). doi:10.32614/CRAN.package.vegan.

53. Dormann, C. F., Fruend, J., Bluethgen, N. & Gruber, B. Indices, graphs and null models: analyzing bipartite ecological networks. The Open Ecology Journal 2, 7–24 (2009).

54. Dormann, C. F., Gruber, B. & Fruend, J. Introducing the bipartite Package: Analysing Ecological Networks. R News 8, 8–11 (2008).

55. PACo: A Novel Procrustes Application to Cophylogenetic Analysis | PLOS One. https://journals.plos.org/plosone/article?id=10.1371/journal.pone.0061048.

56. Kuznetsova, A., Brockhoff, P. B. & Christensen, R. H. B. lmerTest Package: Tests in Linear Mixed Effects Models. Journal of Statistical Software 82, 1–26 (2017).

57. Brooks, M. E. et al. glmmTMB Balances Speed and Flexibility Among Packages for Zero-inflated Generalized Linear Mixed Modeling. The R Journal 9, 378–400 (2017).

58. McGillycuddy, M., Warton, D. I., Popovic, G. & Bolker, B. M. Parsimoniously Fitting Large Multivariate Random Effects in glmmTMB. Journal of Statistical Software 112, 1–19 (2025).

59. Hartig, F. DHARMa: Residual Diagnostics for Hierarchical (Multi-Level / Mixed) Regression Models. (2024). doi:10.32614/CRAN.package.DHARMa.

60. Schmidtke, D. T. et al. The prototypic crAssphage is a linear phage-plasmid. Cell Host Microbe 33, 1347–1362.e5 (2025).

61. Weitz, J. S. et al. Phage–bacteria infection networks. Trends in Microbiology 21, 82–91 (2013).

62. Balbuena, J. A., Míguez-Lozano, R. & Blasco-Costa, I. PACo: A Novel Procrustes Application to Cophylogenetic Analysis. PLOS ONE 8, e61048 (2013).

63. Baykov, I. K. et al. Tentaclins—A Novel Family of Phage Receptor-Binding Proteins That Can Be Hypermutated by DGR Systems. International Journal of Molecular Sciences 24, 17324 (2023).

64. Flores, C. O., Meyer, J. R., Valverde, S., Farr, L. & Weitz, J. S. Statistical structure of host–phage interactions. Proc Natl Acad Sci U S A 108, E288–E297 (2011).

65. Beckett, S. J. & Williams, H. T. P. Coevolutionary diversification creates nested-modular structure in phage–bacteria interaction networks. Interface Focus 3, 20130033 (2013).

66. Gómez, P. & Buckling, A. Bacteria-Phage Antagonistic Coevolution in Soil. Science 332, 106–109 (2011).

67. Wendling, C. C., Goehlich, H. & Roth, O. The structure of temperate phage–bacteria infection networks changes with the phylogenetic distance of the host bacteria. Biology Letters 14, 20180320 (2018).

